# RNA-to-image multi-cancer synthesis using cascaded diffusion models

**DOI:** 10.1101/2023.01.13.523899

**Authors:** Francisco Carrillo-Perez, Marija Pizurica, Yuanning Zheng, Tarak Nath Nandi, Ravi Madduri, Jeanne Shen, Olivier Gevaert

## Abstract

Data scarcity presents a significant obstacle in the field of biomedicine, where acquiring diverse and sufficient datasets can be costly and challenging. Synthetic data generation offers a potential solution to this problem by expanding dataset sizes, thereby enabling the training of more robust and generalizable machine learning models. Although previous studies have explored synthetic data generation for cancer diagnosis, they have predominantly focused on single modality settings, such as whole-slide image tiles or RNA-Seq data. To bridge this gap, we propose a novel approach, RNA-Cascaded-Diffusion-Model or RNA-CDM, for performing RNA-to-image synthesis in a multi-cancer context, drawing inspiration from successful text-to-image synthesis models used in natural images. In our approach, we employ a variational auto-encoder to reduce the dimensionality of a patient’s gene expression profile, effectively distinguishing between different types of cancer. Subsequently, we employ a cascaded diffusion model to synthesize realistic whole-slide image tiles using the latent representation derived from the patient’s RNA-Seq data. Our results demonstrate that the generated tiles accurately preserve the distribution of cell types observed in real-world data, with state-of-the-art cell identification models successfully detecting important cell types in the synthetic samples. Furthermore, we illustrate that the synthetic tiles maintain the cell fraction observed in bulk RNA-Seq data and that modifications in gene expression affect the composition of cell types in the synthetic tiles. Next, we utilize the synthetic data generated by RNA-CDM to pretrain machine learning models and observe improved performance compared to training from scratch. Our study emphasizes the potential usefulness of synthetic data in developing machine learning models in sarce-data settings, while also highlighting the possibility of imputing missing data modalities by leveraging the available information. In conclusion, our proposed RNA-CDM approach for synthetic data generation in biomedicine, particularly in the context of cancer diagnosis, offers a novel and promising solution to address data scarcity. By generating synthetic data that aligns with real-world distributions and leveraging it to pretrain machine learning models, we contribute to the development of robust clinical decision support systems and potential advancements in precision medicine.

## Introduction

Cancer is one of the leading causes of death worldwide, just behind cardiovascular diseases [1]. A physician usually carries out multiple screenings to diagnose the disease in a clinical setting, such as visually examining a digitized tissue slide or finding specific up or down regulations in the patient’s gene expression. Even though these screenings are routinely used at hospitals, rarely all of them are performed on the same patient due to monetary or logistical constraints. Cancer is a multi-scale and multi-factorial disease, and its effects can be recognized at multiple levels [2–4]. For instance, genetic alterations in tumor cells and cells from the tumor microenvironment lead to functional changes which in turn can influence their cellular physiology. [5–8]. Therefore, by not having all screening modalities available, we are losing a part of the picture that can lead to early detection.

Machine learning (ML), and specifically deep learning (DL), has shown tremendous potential for cancer detection and classification in recent years. By using different modalities, such as RNA-Seq, whole-slide-imaging (WSI), miRNA-Seq, or DNA methylation, promising clinical decision support systems have been created [8–12]. However, two problems are present when working with cancer data. First, DL models are known for being data hungry, requiring huge amounts of data to be properly trained. Second, even though the combination of biological data types in a multimodal setting has shown to be superior for cancer detection and prognosis [13–19], unfortunately, the majority of available datasets are incomplete, missing some modalities. Although certain projects make efforts to get a complete picture of the disease by collecting all modalities, such as The Cancer Genome Atlas (TCGA) [20] project or the UK Genomics Pathology Imaging Collection [21], the number of single-modality datasets available greatly surpass them. For instance, there are thousands of gene expression series in the Gene Expression Omnibus (GEO) platform [22] for which the corresponding tissue slide is not available, limiting the potential for creating multimodal ML models.

To address these issues, generative models have been presented as a solution in the literature for cancer data. Specifically, generative adversarial networks (GANs) and Variational Autoencoders (VAE) have been used for generating synthetic WSI and RNA-Seq data. GANs have shown their abilities for modeling cancer characteristics across multiple cancer types, successfully generating synthetic tiles [23, 24] and synthetic gene expression profiles that closely resemble real profiles and capture biological information [25]. VAEs have been successfully applied to gene expression data, showing synthetic generation capabilities in a temporal way and have also been used for data imputation [26, 27]. However, these models present some significant drawbacks when dealing with image data. While sampling is fast for GANs and although they can generate high-quality data, their training is unstable and prone to model collapse, leading to loss of diversity in the generated samples [28–30]. Given these problems, usually a different model is trained for each tissue per cancer type, which is not practical in a real-world scenario. In the case of VAEs, even though their training is much more stable, the generated samples are often blurry compared to those coming from GANs because of the injected noise and the imperfect reconstruction [31]. Furthermore, although models have been presented for the synthetic generation of both WSI and gene expression data, the multi-scale nature of cancer is usually not taken into account for the generation. Specifically, to the best of our knowledge, RNA-to-image synthesis has not been yet explored for cancer tissue.

Text-to-image models have recently caught the attention of the world, given their incredible generative performance. Dall-E 2 and Imagen presented incredible results in image synthesis based on textual input, surpassing all previous methods in quality metrics and synthesis capabilities [32, 33]. They rely on a novel paradigm that is different from previous GAN and VAE-based approaches. Instead, they make use of diffusion models, a DL technique firstly introduced by Sohl-Dickstein et al. [34]. Diffusion models are based on Langevin dynamics, which is an approach in physics for the mathematical modelling of the dynamics of molecular systems. Random noise is incrementally added to the data, thereby ‘destroying’ it until an isotropic random Gaussian distribution is left. Then, a model is trained that learns to reverse this process. Once the model is trained, new images can be synthesized from noise, yet resemble the images from the training data. Different approaches are used for conditioning the model to generate different images based on the given text. Authors of Imagen used a pre-trained large language model to obtain an embedding of the text to condition or ‘guide’ the model generation. Similarly, Dall-E 2 authors obtained a CLIP image-text embedding [35] to condition the diffusion model during the generation step. New images are created using these two approaches, capturing the context of the text and portraying it in the generated image.

Several works have been presented in literature assessing the role of gene expression and histology [36–38], showing that morphological characteristics present in histology associate to changes in gene expression. Inspired by these and the rise of text-to-image models, we explore the relation between cancer tissue and its gene expression profile in an RNA-to-image synthesis problem, with the goal of using synthetic cancer images to pretrain DL models and impute missing data modalities. We present RNA-CDM, a single-cascaded diffusion-based model that is able to synthesize routine hematoxylin and eosin (H&E)-stained tissue image tiles for different cancer tissues without specifying the tissue label, by conditioning on a latent representation of the gene expression profile obtained using a *β*-VAE.

Using a state-of-the-art cell segmentation model, we show that the generated tiles maintain the cell distribution of the real data, preserving cell morphology and specific cell-fractions in the bulk RNA-Seq. We further show that changes in gene expression markers of specific cell types (e.g. lymphocytes) affect the prevalence of those cell types in generated tiles. Finally, we prove that the synthetic data can be used for pretraining models which boost the performance on biomedical classification tasks, thereby showing the potential of generated synthetic tiles for both boosting the pretraining DL models and imputing missing modalities.

## Results

### RNA-CDM can perform realistic RNA-to-image multi-cancer synthesis

Given the high dimensionality of the RNA-Seq data, directly using it for conditioning the diffusion process is not possible. Therefore, inspired by text-to-image synthesis work, we trained a *β*VAE to project the RNA-Seq data from twelve different cancer tissues into a lower-dimensional latent space (Table 2). We decided to use a *β*VAE, given its superior performance in the literature, particularly in comparison to standard autoencoders [27]. The *β*VAE obtained a root mean squared error of 0.2475 and a mean absolute error of 0.1471 in the test set (see Methods for details). We visualized the reconstructed RNA-Seq data using the Uniform Manifold Approximation and Projection (UMAP) algorithm [39] showing different clusters for each cancer tissue (Figure 2).

**Fig. 1.**
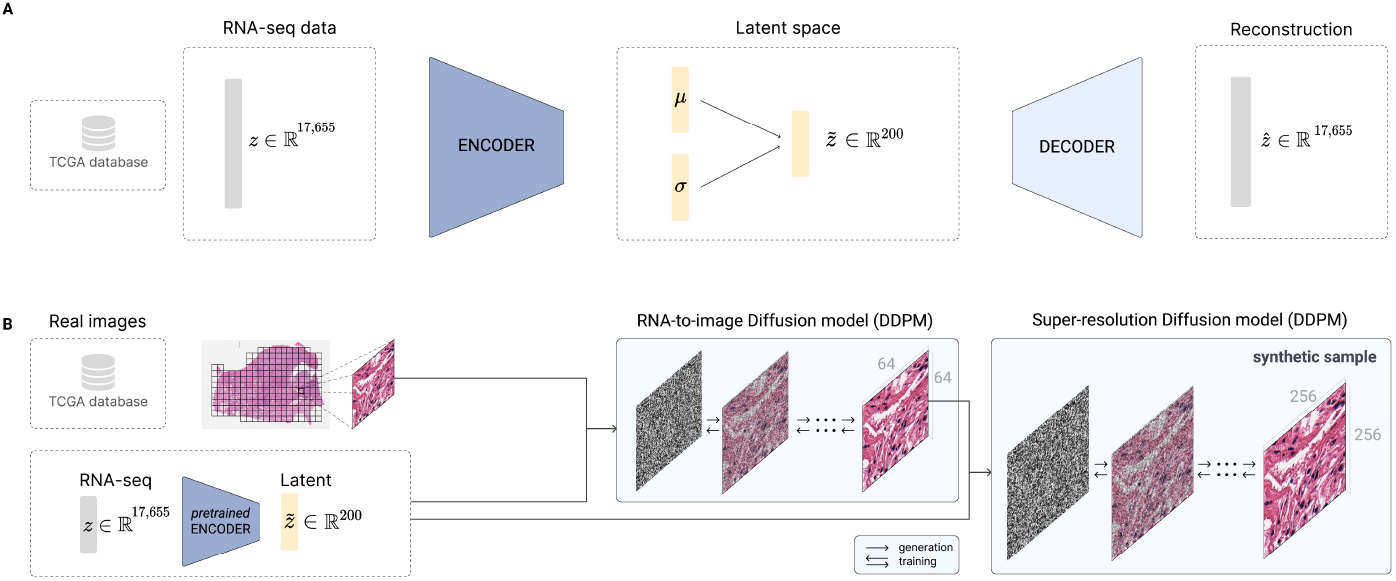
RNA-CDM model architecture used for the generation of RNA-Seq embeddings and synthetic WSI tiles using diffusion models. Panel A: *β*VAE architecture for the generation of gene expression embeddings. The model uses as input the expression of 17, 655 genes. Both the encoder and the decoder are formed by two linear layers of 6, 000 and 4000 respectively. The latent *µ* and *σ* vectors have a feature size of 200. **Panel B:** RNA-CDM architecture for the generation of synthetic multi-cancer tiles. It is formed by two Denoising Diffusion Probabilistic Models (DDPM), one acting as a RNA-to-image model and the second one as a super-resolution model. The pre-trained encoder is used to obtain the latent representation of the RNA-Seq data. A corresponding tile from the patient is obtained. For the first DDPM, the tile is resized from 256×256 to 64×64 pixels. During the training phase, noise is gradually applied to the tile according to the given noise scheduler at each timestep *t*. Then, the first DDPM learns to reduce the noise by using as input the noisy image, the timestep *t*, and the gene expression embedding. The noise predicted is removed from the noisy image at each time step, having a denoised tile of 64×64 at the end of the process. Then, a second DDPM takes the denoised image, the noisy 256×256 image at timestep *t*, the timestep *t*, and the gene expression embedding and predicts the added noise again. Then, the noise is removed from the 256×256 tile iteratively until an denoised image is obtained and compared with the original tile. For generating a new image, the process is the same, but we start from total random noise until we have a synthetic tile whose generation has been guided by the gene expression embedding.

Next, we trained the RNA-CDM model using multimodal data from five cancer tissues: lung adenocarcinoma (LUAD), kidney renal papillary cell carcinoma (KIRP), cervical squamous cell carcinoma (CESC), colon adenocarcinoma (COAD) and glioblastoma (GBM). Our RNA-CDM model was able to accurately generate tiles from the five cancer types without any tissue label information and only conditioning on the latent representation of the RNA-Seq data (Figure 2). Various tissue morphology can be observed in the generated tiles, from more homogeneous and cell-abundant (e.g. all the different types of glial cells) GBM tissue to the more sparse LUAD tissue. For all generated tiles, cell nuclei and muscle fibers can be distinguished in the tiles (Supplementary Figure 1).

**Fig. 2.**
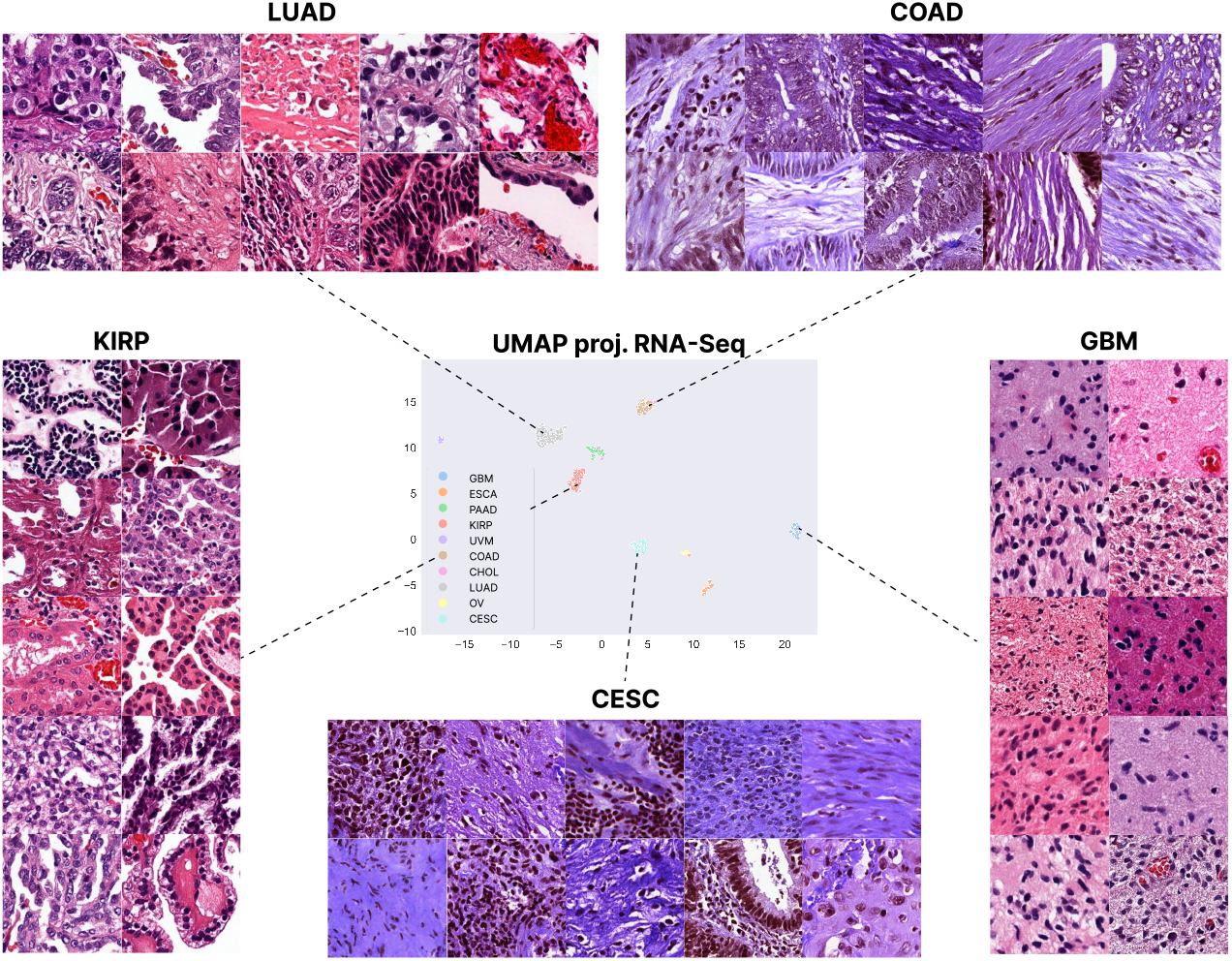
RNA-to-image multi-cancer synthetic samples generated by conditioning on the gene expression latent representation. The *β*VAE is able to obtain an accurate representation of the multi-cancer gene expression profiles. Once this model has been trained, the RNA-CDM architecture (see Figure 1 and Subsection 6) is trained and conditioned on the latent representation of the RNA-Seq data over five different cancer types: lung adenocarcinoma (LUAD), kidney renal papillary cell carcinoma (KIRP), cervical squamous cell carcinoma (CESC), colon adenocarcinoma (COAD) and glioblastoma (GBM)). The model accurately synthesizes the samples, capturing the distinct morphologic characteristics of each cancer type.

We evaluated the in-silico quality of the tiles using the standard quality evaluation metrics for generative models, such as the Frechet Inception Distance (FID) [40], the Inception Score (IS) [30], and the Kernel Inception Distance (KID) [41]. Hereto, we generated 50*k* tiles (10, 000 per cancer type), and compared them with the same amount of real tiles. The generated tiles obtained a FID50k score of 23.36, an IS50k of 3.19, and a KID50k of 0.015.

To validate the generalization capabilities of the model, we obtained RNA-Seq data from two additional external data sets: a colorectal RNA-Seq data set (accession number: GSE50760 [42]) and a lung cancer RNA-Seq data set (accession number: GSE226069 [43]). RNA-CDM was able to accurately generate synthetic tiles for both cancer types, showing its generalization capabilities (Supplementary Figure 2).

### RNA-CDM tiles maintain the cell distribution of real tiles and significantly correlate with deconvolved cell fractions

Next, we used HoverNet [44], a state-of-the-art cell segmentation and classification model, over real and synthetic tiles to detect different cell types. We found that the distribution of cells in real and synthetic tiles was similar across the five cancer tissues (Figure 3A), and the mean number of cells detected across cell-types was similar for the majority of cancer and cell types (Table 1). Even though HoverNet was only trained on real data, cell types were correctly detected in the synthetic tiles, showing that RNA-CDM can produce samples with realistic cell morphology (Figure 3B).

**Fig. 3.**
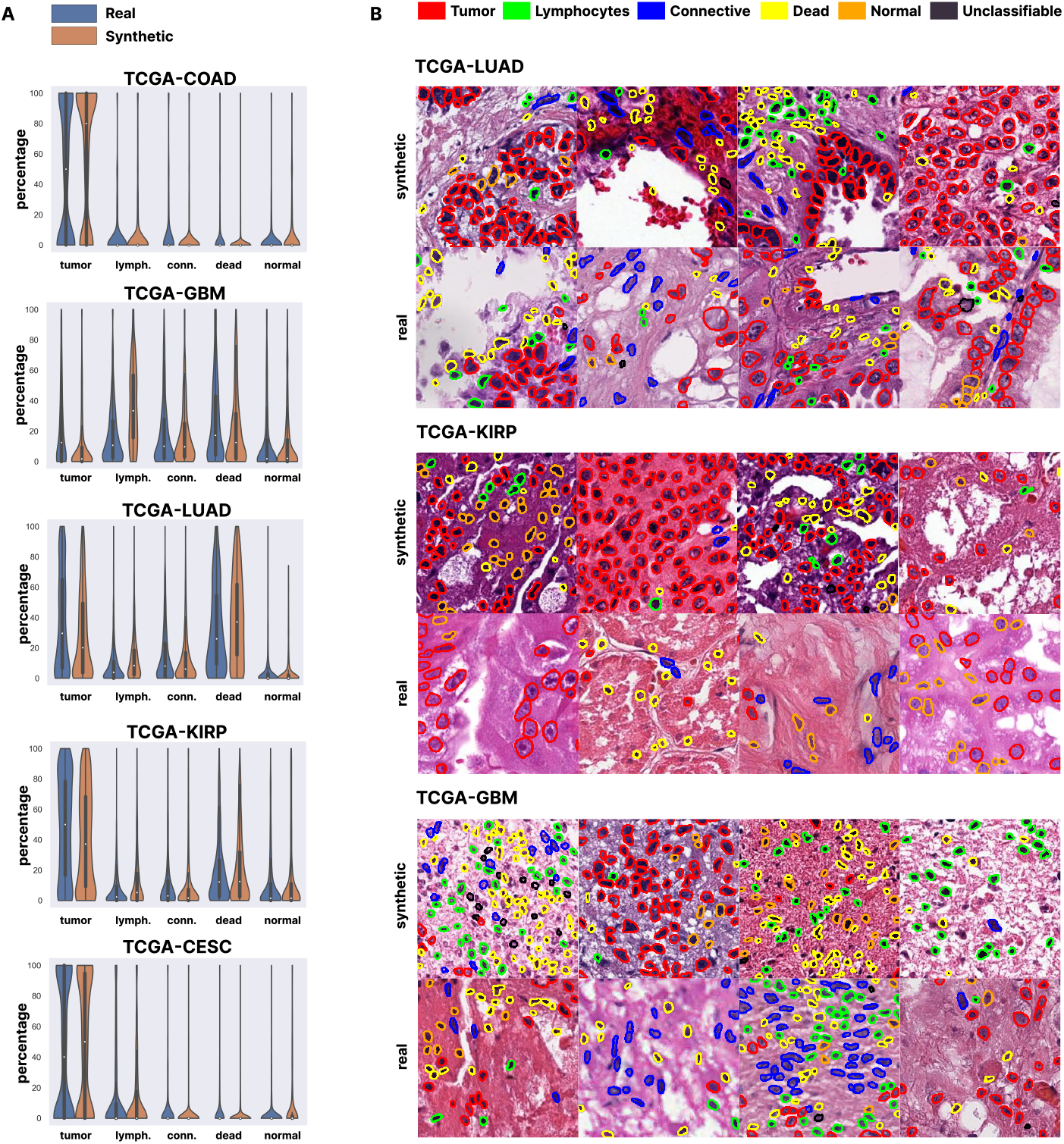
Synthetic samples maintain the cell distributions observed in real world data. **Panel A:** Distribution of cell types is maintained across the different cancer types in both real and synthetic tiles generated using RNA-CDM. We used HoverNet pretrained weights on the PanNuke dataset to detect five different cell types (tumor, lymphocytes, connective, necrotic, and normal). Normal in the PanNuke dataset includes cells from normal to degenerative, metaplastic, atypia etc. 100, 000 tiles were used in total (50, 000 real and 50, 000 synthetic, with 10, 000 tiles per cancer type class). **Panel B:** Examples of cells detected in real and synthetic tiles by HoverNet. Cells are detected in both cases, showing that cell morphology is maintained in the RNA-CDM generated tiles.

**Table 1.** Mean percentage of cells detected by HoverNet on real and RNA-CDM generated tiles for each cancer type.

|  | Tile type | Tumor | Lymphocytes | Connective | Dead | Normal |
| --- | --- | --- | --- | --- | --- | --- |
| TCGA-COAD | Real | 47.44 ± 43.12 | 7.69 ± 20.43 | 10.48 ± 24.58 | 4.21 ± 14.66 | 3.94 ± 14.51 |
| | Synthetic | 61.78 ± 42.43 | 6.11 ± 17.88 | 2.69 ± 13.27 | 1.33 ± 7.36 | 6.87 ± 17.89 |
| TCGA-GBM | Real | 22.57 ± 25.83 | 17.66 ± 20.25 | 18.54 ± 21.74 | 26.20 ± 26.20 | 12.50 ± 21.34 |
| | Synthetic | 9.18 ± 16.15 | 35.09 ± 24.89 | 17.11 ± 20.33 | 22.89 ± 24.57 | 11.99 ± 19.76 |
| TCGA-LUAD | Real | 37.40 ± 32.49 | 8.12 ± 11.72 | 15.36 ± 19.22 | 33.15 ± 28.07 | 3.65 ± 9.78 |
| | Synthetic | 27.74 ± 28.62 | 12.93 ± 15.15 | 11.00 ± 15.19 | 41.30 ± 28.98 | 3.31 ± 9.86 |
| TCGA-KIRP | Real | 48.14 ± 33.02 | 7.39 ± 12.11 | 12.57 ± 21.83 | 19.20 ± 21.75 | 10.37 ± 18.35 |
| | Synthetic | 40.85 ± 32.25 | 12.92 ± 17.84 | 8.17 ± 17.38 | 20.59 ± 23.95 | 12.34 ± 21.84 |
| TCGA-CESC | Real | 45.52 ± 43.65 | 13.61 ± 27.54 | 5.34 ± 17.52 | 4.59 ± 15.31 | 2.85 ± 10.87 |
| | Synthetic | 45.82 ± 42.78 | 15.48 ± 28.17 | 1.52 ± 10.29 | 1.70 ± 7.74 | 7.89 ± 19.97 |

We conducted additional tests to determine whether the gene expression profile characteristics remained consistent in the synthetic tiles. Specifically, we examined whether the proportions of different cell types in the bulk RNA-Seq correlated with the cell types found by HoverNet in the synthetic tiles. In all cases cell-fraction percentage significantly correlated with the percentage of corresponding cell types found in the tissue by Hovernet (p-value ≤ 0.05), showing that cell-specific characteristics are preserved in the synthetic tiles generated using the bulk RNA-Seq (Supplementary Figure 3).

Next, we examined whether the use of de-convolved gene expression for generating tiles affects the presence of certain cell types in the synthetic tiles, even though RNA-CDM was not trained with de-convolved gene expression. We tested the generation of synthetic tiles using the fibroblast and haematopoietic de-convolved RNA-Seq. For fibroblast gene expression, we expected to see an increase in the percentage of connective tissue cells in the tiles, likewise for haematopoietic gene expression and the percentage of lymphocytes. We indeed found that the mean percentage of connective tissue cells was higher in tiles generated using the fibroblast RNA-Seq than in tiles generated with bulk RNA-Seq data across all cancer types (Supplementary Figure 4A). Similarly, the mean percentage of lymphocytes is higher when using the haematopoietic deconvolved RNA-Seq in LUAD and KIRP, approximately the same percentage is obtained in COAD and CESC, and a lower value is obtained for GBM, tumors that typically don’t contain lymphocytes [45, 46] (Supplementary Figure 4B).

### RNA-CDM synthetic samples can be used as pretraining to improve classification performance

Next, we experimented with the utility of synthetic data for training DL models in settings of varying availability of real training data. To do so, we used 5, 000 tiles, consisting either of 100% real tiles or of which a certain percentage (25%, 50%, 75%) is replaced with synthetic tiles (simulating situations with fewer available real data). We evaluated the performance of classifying each tile into one of five different tissue classes using a ResNet18 model and 5-fold cross validation (5-Fold CV). In all cases, there was no difference in the classification performance, showing that synthetic data can accurately substitute real data with no effect on the classification task.

Next, we also experimented with using exclusively synthetic data for pretraining a classification model. We used four different sizes for the pretraining dataset 25% (1,250 samples), 50% (2,500 samples), 75% (3,750 samples) and 100% (5,000 samples) and compared how this affected the classification performance on a real sample dataset.

We found that using the synthetic data for pretraining improved the model’s performance over the baseline, regardless of the size of the pretraining dataset, for both the accuracy and F1-score (see Figure 4 B). The performance of the model also increased with the size of the synthetic dataset, showing that pretraining with more synthetic samples boosted the model’s performance on the real dataset. To further quantify the observed improvement, we calculated confusion matrices over all of the real samples for the model with and without pretraining (Figure 4 C).

**Fig. 4.**
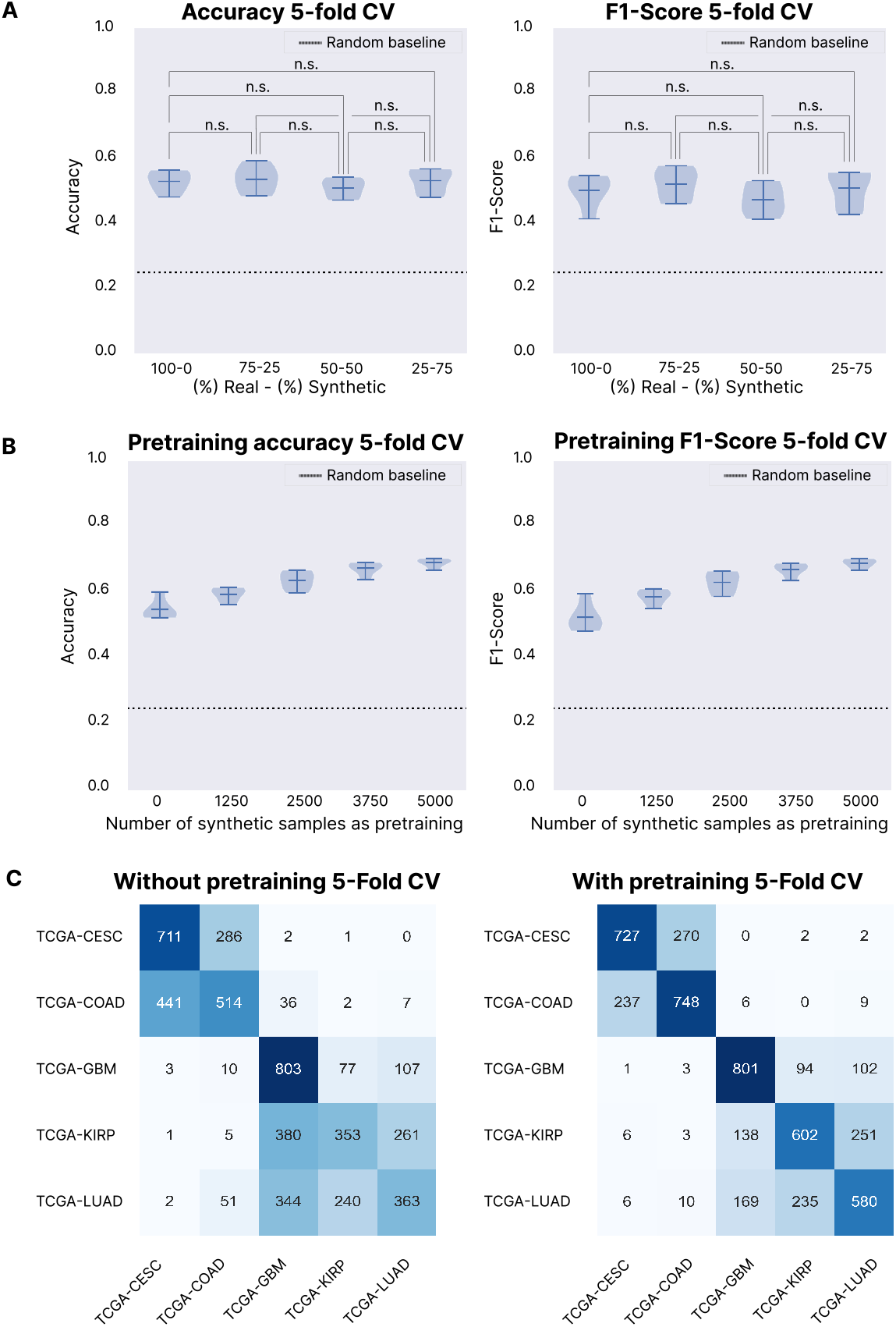
Pretraining on synthetic samples improves classification performance in a multi-cancer classification problem. **Panel A:** we substituted different percentages of the training data with synthetic samples generated by RNA-CDM in a 5-Fold CV over the real data (classification between the five different cancer types). The performance does not decrease significantly in any experimental setting (n.s. states for not significant, p-value ≤ 0.05.) **Panel B:** we used different numbers of synthetic data samples as pretraining dataset. In both accuracy and F1-Score, increasing the number of samples in the pretraining dataset increases the performance in a 5-Fold CV over the real data. **Panel C:** confusion matrix of all the test sets in the 5-Fold CV without using any pretraining. (On the right) Confusion matrix of all the test sets in the 5-Fold CV using all the synthetic samples as pretraining. The model with the pretrained weights improves the classification performance and does not introduce any bias or new errors between the classes.

To show the added value of synthetic tiles for model training, we approached the problem of microsatellite instability status prediction in colorectal cancer, comparing a model pretrained on synthetic tiles using self-supervised learning (SSL) and a model trained from scratch. The model using the SSL pre-trained weighs outperforms the model trained from scratch in all four cases, showing a difference in accuracy of up to 11.8 percentage when both models are trained with 1000 samples (Supplementary Figure 5). The best performance was obtained with the model using the SSL weights and trained over 4000 samples, reaching a mean accuracy of 73.42 ± 1.5. The maximum accuracy of the model trained from scratch is also obtained when trained over 4000 samples, but only reaches an accuracy of 66.06 ± 2.1, only surpassing the performance of the SSL weights model when trained on 200 samples.

## Discussion

Association of molecular and morphologic data of cancer tissues is emerging as an important research area in computational pathology and oncology [47]. Quantitative models have been developed to predict molecular biomarker status directly from H&E-stained WSI (including prediction of microsatellite instability for colorectal cancer [48] and EGFR mutations for lung cancer [8]) opening an exciting avenue for the development of “digital biomarker” tests which might obviate the need for more expensive and time-consuming assays. However, a significant barrier to progress in the development of such tests is the relative paucity of tumor H&E specimens with corresponding ground truth labels availability for model training. While most patients have H&E tumor specimens collected as part of routine cancer diagnostic workflows, far fewer patients receive biomarker testing on those specimens, and when such biomarker tests are performed, an accompanying H&E-stained slide corresponding to the test sample might not be generated (for example, if the entire sample is exhausted during performance of the molecular assay). The ability of the RNA-CDM model in this study to generate synthetic but realistic H&E training images only from RNA-Seq expression data, and the demonstration that the resultant synthetic training data can be used to improve the performance of a downstream tumor classification model, offers a promising solution to the data scarcity problem.

While most models have focused on the detection of either discrete molecular changes, such as mutations in a specific gene or expression of a single protein, or classification of tumors into a limited number of discrete classes such as consensus molecular subtypes [49], few attempts have been made to explore how changes in the expression of thousands of genes might be reflected in routine H&E tumor histomorphology. Although spatial profiling methods (such as Akoya, 10x Genomics, etc) allow for some correlation between gene expression profiles and histology, these methods are expensive, currently impractical on a large scale and rely mainly on identification of pre-determined/handcrafted features or cell types that are recognizable to the human eye. Therefore, biologically important sub-visual morphologic features present on slides might be missed. Aside from their practical utility in data augmentation, models such as the RNA-CDM model developed in this study, which utilize latent representations of an entire RNA-Seq profile, might allow for the identification of novel morphologic features associated with clinically relevant molecular biological states that are currently unrecognized by the human eye.

Previous approaches for WSI cancer tile generation have primarily relied on GAN architectures and were able to generate realistic-appearing tissue images [23, 24]. However, GANs are prone to mode collapse and generate a low diversity of samples, limiting their potential for synthetic data generation that reflects real-world data distributions [28–30]. Unlike natural images, cancer tissues are generally more homogeneous, requiring training of a different model for each cancer type in order to avoid mode collapse to a single class, which makes GANs impractical for generating cancer images in real-world scenarios [23, 24, 50]. Furthermore, previous approaches for generating synthetic WSI have not made use of RNA expression data. Here, we present RNA-CDM, a single architecture based on cascaded diffusion models capable of performing RNA-to-image multi-cancer synthesis without any explicit label information (e.g. the cancer type), just using the expression profile of the patient. Because RNA-CDM uses only the latent representation obtained using the *β*VAE model (Figure 1 A) and requires only a single architecture to generate data for five different cancer types, it is more efficient and practical for synthetic data generation. This means that multiple models do not need to be implemented, saving on computational and storage costs. However, it must be mentioned that when using the Karras et al. [51] sampling method, the sampling time is faster for GANs. We expect that future research will address this drawback of diffusion models. Previous models, such as those presented by Quiros et al. [23], obtained an FID score of 16.65 on breast cancer and 32.05 on colorectal cancer. Our model obtains similar results with an FID score of 31.70 for colorectal cancer. In addition, our proposed RNA-CDM model generates tiles from five different cancer types using a single architecture, while Quiros et al. trained a dedicated model per cancer type. Training an RNA-CDM on a single cancer likely can improve the FID score significantly.

Next, we demonstrated the consistency of cell distribution patterns between synthetic and real tiles across various cancer types (Figure 3A, Table 1). Additionally, we observed persistent differences in cell composition among different cancer types (Figure 3A). Notably, cell types that are more prevalent in specific cancer types, such as dead cells in LUAD, also exhibit higher abundance in the synthetic tiles (Table 1). Similarly, normal cells are more abundant in both GBM and KIRP real and synthetic tiles, in comparison to the rest of cancer types. Tumor cells are the most predominant cell type found both in real and synthetic tiles, with the lowest amount in GBM. These findings demonstrate the model’s capability to capture cancer-specific cell characteristics, and that real tiles’ trends are maintained in the synthetic tiles.

To further validate these results, we performed deconvolution of bulk RNA-Seq data and established a significant correlation between the percentage of specific cell types identified in the generated tiles by HoverNet and the corresponding cell fraction predicted by CIBERSORTx (Supplementary Figure 3). These findings highlight the importance of obtaining the latent representation of gene expression and the model’s ability to capture latent information from the entire gene set during training, which would not have been possible when conditioning only on a simple cancer-level label.

Furthermore, we investigated the impact of de-convolved data by generating synthetic tiles using fibroblast and hematopoietic RNA-Seq. Using these two input types resulted in respectively the identification of a higher number of connective tissue cells and lymphocytes in the synthetic tiles (Supplementary Figure 4), demonstrating the influence of the latent gene expression representation. Our findings based on the de-convolved hematopoietic data showed that the lymphocyte proportion was lower in GBM compared to the other cancer types. This result is in agreement with previous studies as GBMs contain a limited number of tumor infiltrating lymphocytes and the majority of immune cells in brain tumors are macrophages [45, 46]. Therefore, although the model was not explicitly trained on de-convolved gene expression, it successfully captured the relationship between specific cell types’ expressionf and their influence on tissue morphology. Further investigations are warranted to explore potential enhancements with more diverse and comprehensive data.

Next, we focused on one of the main utilities of synthetic data, increasing the size of small datasets to boost the performance of machine learning models [52]. We first tested whether substituting real data with synthetic data impacted the performance of a multi-cancer classification model. For all experimental settings, no significant different was found in the classification performance, highlighting the quality of the synthetic tiles (Figure 4 A). We also evaluated the benefit of using synthetic data for model pretraining. Both accuracy and F1-Score increased, irrespective of the size of the pretraining dataset (Figure 4 B). These results are promising and show that increasing the number of synthetic samples in the pretraining set directly improves the classification performance. The classification confusion matrices also showed that the fine-tuned model improves the model trained from scratch while maintaining the class-specific misclassifications, so we are not introducing new biases or errors with the synthetic data (Figure 4). Furthermore, we investigated the application of SSL to enhance the classification performance of our model in a biologically relevant task, namely, microsatellite instability status prediction. By leveraging SSL with learned weights, we demonstrated improved classification performance compared to training a model from scratch (Supplementary Figure 5). Our synthetically generated tiles, serving as unlabeled realistic data, thus can facilitate the learning of general patterns that can be subsequently fine-tuned for downstream tasks. The effectiveness of SSL techniques has previously been demonstrated in various medically-related domains [12, 53], highlighting their potential for leveraging unlabeled data. In this context, RNA-CDM serves as a valuable data generator for enhancing the performance of SSL models in biomedical downstream tasks.

One limitation of this study is that we are using TCGA to train our model. TCGA is the largest data set for cancer research and the tissue samples are reviewed by a pathologist to confirm the diagnosis and that the sample meets inclusion criteria, more specifically samples need to contain at least 60% tumor nuclei and have less than 20% necrotic tissue.

In summary, we have proposed a solution for data scarcity in machine learning problems by using an RNA-to-image multi-cancer cascaded diffusion model. We show how changes in gene expression can be studied in-silico by using RNA-CDM, exploring the interactions between the two modalities without the need of generating new data and improving classification problems. New technologies such as spatial transcriptomics [54] are emerging that generate a spatial map of gene expression across the tissue. We expect that these spatial technologies will further enhance the capabilities of models such as RNA-CDM.

## Materials and Methods

### Data acquisition and data pre-processing

Data were obtained from the TCGA project database, which contains paired samples of RNA-Seq and WSIs from patients. For the WSI data, only diagnostic slides were considered, given the superior histologic quality of the sample. For the gene expression data, we downloaded the raw files for later preprocessing steps. Twelve cancer types were considered for the training of the *β*-VAE, while five were used during the training and validation of RNA-CDM (TCGA-LUAD, TCGA-GBM, TCGA-KIRP, TCGA-CESC, TCGA-COAD). These cancer types were selected based on their morphology differences and the number of samples available, given the data requirements of the model training. The cancer types used and the number of available samples are presented in Table 2.

**Table 2.**
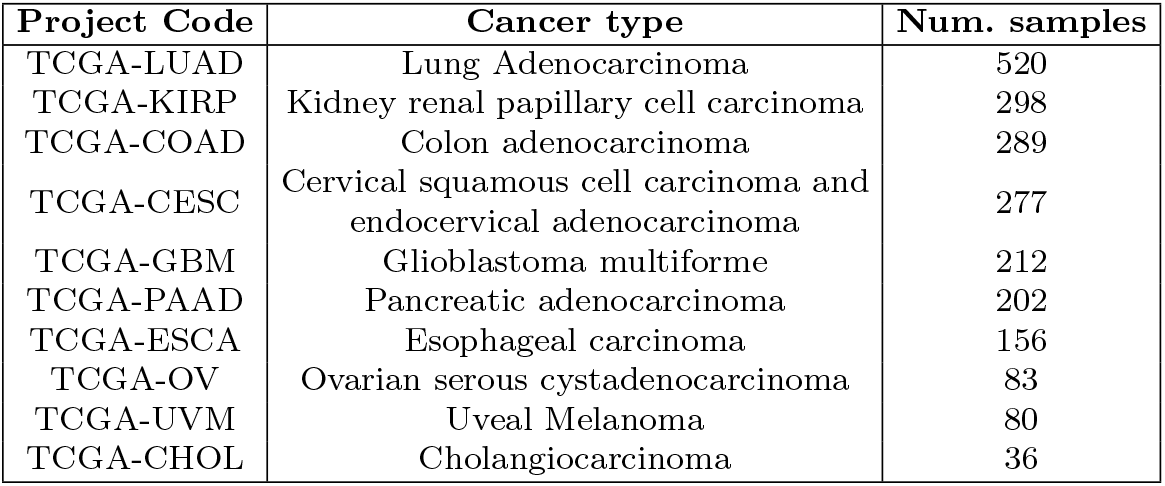
TCGA projects used for the training of the *β*-VAE and the RNA-CDM model.

| Project Code | Cancer type | Num. samples |
| --- | --- | --- |
| TCGA-LUAD | Lung Adenocarcinoma | 520 |
| TCGA-KIRP | Kidney renal papillary cell carcinoma | 298 |
| TCGA-COAD | Colon adenocarcinoma | 289 |
| TCGA-CESC | Cervical squamous cell carcinoma and endocervical adenocarcinoma | 277 |
| TCGA-GBM | Glioblastoma multiforme | 212 |
| TCGA-PAAD | Pancreatic adenocarcinoma | 202 |
| TCGA-ESCA | Esophageal carcinoma | 156 |
| TCGA-OV | Ovarian serous cystadenocarcinoma | 83 |
| TCGA-UVM | Uveal Melanoma | 80 |
| TCGA-CHOL | Cholangiocarcinoma | 36 |

Once the data were downloaded, we proceeded with the pre-processing steps. For the gene expression data, we followed the pre-processing steps described by Zheng et al. [55]. The raw sequencing reads were aligned to the human transcriptome and quantified using the Kallisto-Sleuth algorithm proposed by Bray et al. [56]. NaN values were removed and we selected those genes the different cancer types have in common. These pre-processing steps left us with a total of 17, 655 genes that were used as input to the model. The gene expression was firstly log-transformed and then normalized with the z-score transformation from the training set values.

WSIs were acquired in SVS format and downsampled to 20× magnification (0.5*µm* px^-1^). The size of WSIs is usually over 10*k ×* 10*k* pixels, and therefore, cannot be directly used for training machine learning models to generate the data. Instead, tiles of a certain dimension are taken from the tissue, and they are used to train the models, which is consistent with related work in state-of-the-art WSI processing [8, 57, 58]. In our work, we took non-overlapping tiles of 256 × 256 pixels. Firstly, a mask of the tissue in the higher resolution of the SVS file was obtained using the Otsu threshold method [59]. Tiles containing more than 60% of the background and with low contrast were discarded. A maximum of 4, 000 tiles were taken from each slide. For the preprocessing of the images we relied on the python package openslide [60], which allows us to efficiently work with WSI images. The tiles were saved in an LMDB database using as an index the number of the tile. This approach enables us to reduce the number of generated files, and structure the tiles in an organized way for a faster reading while training. Tiles containing pen marks or other artifacts were filtered during the reading phase.

For the generalization experiments, two series were downloaded from GEO, GSE50760 [42]) and GSE226069 [43]. Since we need to first obtain the latent representation of the RNA-Seq, those genes in common with TCGA were extracted, and those not present in the GEO series were set to 0. Then, data was log-transformed and normalized using z-score based on the TCGA data values.

Data for the microsatellite instability status prediction was obtained from Kather et al. [61], and downloaded from the Kaggle platform ^1^. The dataset contains the patches corresponding to MSS and MSI labels. 75, 039 belonged to the MSI class and 117, 273 to the MSS class.

### *β*-VAE for multi-cancer latent embedding generation

We chose the *β*-VAE model for the creation of a latent embedding [62]. The *β*-VAE model is an extension of the VAE where a *β* parameter is introduced in the loss function. VAEs are an modification of the original autoencoder, but they are both formed by two networks, the encoder and the decoder. The idea behind the autoencoder is to learn a smaller representation of the input data by learning the function *h_θ_*(*x*) ≈ *x* being *θ* the parameters of the neural network. We obtain a lower dimensional representation of the data, and then we reconstruct the input using the decoder. Thus, we want to minimize the reconstruction error between the input and the output. On the other hand, the VAE extends this approach to learn a probability distribution of the latent space. The assumption of the VAE is that the distribution of the data *x*, *P* (*x*) is related to the distribution of the latent variable *z*, *P* (*z*). The loss function of the VAE, which is the negative log-likelihood with a regularizer is as follows:

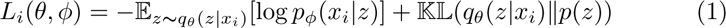

where the first term is the reconstruction loss and the second term is the Kullback-Leibler (KL) divergence between the encoder’s distribution *q_θ_*(*z|x*) and *p*(*z*) which is defined as the standard normal distribution *p*(*z*) = *N* (0, 1).

For the *β*VAE we introduce the parameter *β*, which controls the effect of the KL divergence part of the equation:

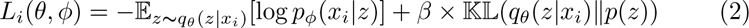

If *β* = 1, we have the standard loss of the VAE. If *β* = 0, we would only focus on the reconstruction loss, approximating the model to a normal autoencoder. For the rest of the values, we are regularizing the effect of the KL divergence on the training of the model, making the latent space smoother and more disentangled [62].

For the final architecture, we empirically determined to use two hidden layers of 6, 000 and 4000 neurons each for both the encoder and the decoder, and size of 200 for the latent dimension, similar to the architecture proposed by Qiu et al. [27]. We used batch norm between the layers and the LeakyReLU as the activation function. A *β* = 0.005 was used in the loss function. We used the Adam optimizer for the training with a learning rate equal to 3 × 10*^−^*^3^, along with a warm-up and a cosine learning rate scheduler and the mean square error as the loss function. We trained the model for 250 epochs with early stopping based on the validation set loss, and a batch size of 128. A schema of the architecture is presented in Figure 1 A. We divided the dataset into 60-20-20 % training, validation and test stratified splits, and we trained the model with twelve different cancer types in order to obtain a better and more general latent space (see Table 2).

### RNA-CDM: A Cascaded diffusion model for multi-cancer RNA-to-image synthesis

We present RNA-CDM, a cascaded diffusion model for multi-cancer RNA-to-image synthesis. Diffusion models are a kind of score-based generative models that model the gradient of the log probability density function using score matching [63, 64]. The idea for diffusion models is to learn a series of state transitions to map noise ɛ from a known prior distribution to *x*_0_ from the data distribution. Firstly, we define an additive noise forward process from *x*_0_ to *x_t_* defined as:

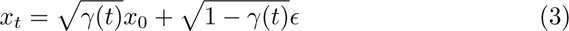

where ɛ ∼ *N* (0, *I*), *t ∼ U* (0, *T*), and *γ*(*t*) is a monotonically decreasing function from 1 to 0. Then, we learn a neural network, *f* (*x_t_, t*) to reverse this process by predicting *x*_0_ (or ɛ) from *x_t_*. The training of the neural network is based on denoising with a *l*_2_ regression loss:

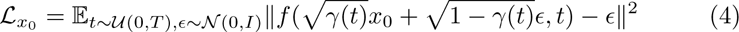

Once we have learned a model, new samples can be generated by reversely going from *x_t_ → x_t−n_ → … → x*_0_. This can be achieved by applying the denoising function *f* to the samples to obtain *x*_0_, and then make the transition to *x_t−n_* by using the predicted *x*_0_ [65].

Cascaded diffusion models were proposed by Ho et al. [66] as a way to improve sample quality. Having high-resolution data *x*_0_ and a low-resolution version *z*_0_, we have a diffusion model at the low resolution *p_θ_*(*z*_0_), and a super-resolution diffusion model *p_θ_*(*x*_0_|*z*_0_). The cascading pipeline forms a latent variable model for high resolution data, that can also be extended to conditioning to the class (or the gene expression latent representation in our case):

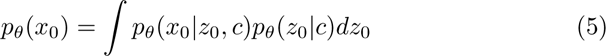

where *c* is the gene expression latent representation. Also, as in normal diffusion models, we condition on the timesteps both for the training and the sampling.

In this work, we used the extension presented by Karras et al. [51], where the training and sampling methods are modified. They proposed a stochastic sampling method, that hugely speeds the sampling process, one of the bottlenecks of diffusion models. More information about their approach can be read in Sections 4 and 5 of their paper [51]. We empirically selected a value of *ρ* = 7, *S_churn_* = 80, *S_tmin_* = 0.05, *S_tmax_* = 50, *S_noise_* = 1.003 and 32 steps for the stochastic sampling.

For the hyperparameters and architectures used, we followed those presented in the Imagen paper [33]. The Unet model [67] was chosen for the diffusion models, using a dimension of 256 for the low-resolution and 128 for the super-resolution diffusion models. Attention and skip connections are used across the Unet layers. The low-resolution diffusion models generate an image of 64×64, and it is conditioned on the RNA-Seq latent representation and the timestep. Then, a gaussian blur is applied to that image and it serves as input to the super-resolution diffusion model (along with the RNA-Seq latent embedding and the timestep), which generates an image of 256×256. The whole architecture accounts for a total of 1.8 billion parameters. A schematic representation of the training and sampling pipeline is presented in Figure 1 B.

### RNA-CDM computational resources and training details

The RNA-CDM model was trained using 4 NVIDIA A100-SXM4-40GB GPUs on the Polaris supercomputer at Argonne National Laboratory. Training was carried out until the visual quality of the generated tissues reached a satisfactory level. In total, the training process required 79*k* steps, where each step involved updating the weights after computing the loss over a batch. A batch size of 64 was utilized per GPU, resulting in an effective batch size of 256. The model was trained for approximately 8 days with this hardware.

Adam optimizer with a learning rate of 1*e^−^*^4^ was used for training. Two versions of the model were maintained during training: i) the training network, that was actually trained using the Adam optimizer, and ii) the EMA network, that stored the exponential moving average (EMA) of the weights of the training network from the previous training steps, was used for sampling. The EMA network owing to its smoother weights obtained from averaging over multiple training steps (instead of using the model parameters from the last training iteration), is less sensitive to local fluctuations during training and hence more amenable for generating realistic synthetic samples as well as less prone to overfitting. For both Unets we used *timesteps* = 1000 with a linear gaussian diffusion process. We used a maximum of randomly chosen 4000 tiles per slide to train the model.

### HoverNet and deconvolution experiments

We used HoverNet [44], a state-of-the-art cell segmentation and classification model, to detect different cell types in cancer tissues. We used the weights trained on the PanNuke dataset to detect the following cell types: tumor, lymphocytes, connective, dead, and normal cells. We generated 50, 000 synthetic samples using multiple RNA-Seq profiles and obtained 50, 000 tiles from real whole-slide images for each of the five tumor tissues. We then compared the distribution of the different cell types between real and synthetic tiles.

We conducted additional tests to determine whether the gene expression profiles characteristics remained consistent in the synthetic tiles. Specifically, we examined whether the proportions of different cell types in the bulk RNA-Seq correlated with the cell types found in the synthetic tiles. We used CIBERSORTx [68, 69] to deconvolve the bulk RNA-Seq from 50 patients (ten per cancer type)in four different cell types: epithelial, fibroblast, endothelial, and haematopoietic cells. We selected these cell-types given that a higher fraction of each of them should affect the percentage of specific cells in the synthetic tiles. We then generated ten tiles per patient using RNA-CDM and the bulk RNA-Seq data from these patients, and employed HoverNet to detect and quantify the cell percentages within each tile. The percentage of fibroblasts detected in the bulk RNA should be correlated to the percentage of cells classified as connective tissue by HoverNet. Similarly, there should be a correlation between bulk-derived fractions of haematopoietic cells with HoverNet-derived fractions of lymphocytes in the tiles. Further, fractions of epithelial cells (bulk derived) should correlate with fractions of tumor cells (HoverNet) and prevalence of endothelial cells (bulk derived) should correspond to prevalence of cells from connective tissue (HoverNet). We calculated the Pearson correlation coefficient between each de-convolved cell type fraction and the corresponding cell type percentages derived in the tiles with HoverNet.

Then, we tested whether using de-convolved RNA-Seq led to a proliferation of specific cell-types. We tested the generation of synthetic tiles using the fibroblast and haematopoietic de-convolved RNA-Seq, and compared with those generated using bulk RNA-Seq, and compared with those generated using bulk RNA-Seq. We generated ten tiles per patient using RNA-CDM and employed HoverNet to detect and quantify the cell percentages within each tile (connective tissue cells in the case of fibroblast and lymphocytes in the case of haematopoietic).

### Training details for ML experiments

For the ML experiments presented in the results section, we used a Resnet-18 architecture trained from scratch. For the first set of experiments, the model was trained for 20 epochs using AdamW optimizer with a learning rate equal to 3*e^−^*^3^. For the second set of experiment, the model was trained firstly on the synthetic data for 50 epochs using an early stopping strategy using the same optimizer and learning rate value. Then, it was fine-tuned in the real data for 20 epochs using a learning rate value of 3*e^−^*^5^.

For the microsatellite instability status prediction, SimCLR was used using a Resnet-18 as a backbone. SimCLR is a contrastive learning method that maximizes the agreement between two different augmented versions of the same image, thereby learning a relevant feature representation of the image [53]. It was trained on 50, 000 synthetic tiles (10, 000 per cancer type) for 50 epochs. Both models used AdamW as the optimizer, but the model trained from scratch used a learning rate of 3*e^−^*^3^, while the model using SSL weights a learning rate of 3*e^−^*^5^. Both models were trained for 100 epochs. Metrics were obtained across the test sets in a 5-Fold CV experimental setting.

## Footnotes

1 https://www.kaggle.com/datasets/joangibert/tcga_coad_msi_mss_jpg

## Acknowledgments

The results published here are in whole or part based upon data generated by the TCGA Research Network: https://www.cancer.gov/tcga. FCP was supported by the Spanish Ministry of Sciences, Innovation and Universities under Projects RTI-2018-101674-B-I00 and PID2021-128317OB-I00, the project from Junta de Andalucia P20-00163 and a Predoctoral scholarship from the Fulbright Spanish Commission. Research reported here was further supported by the National Cancer Institute (NCI) under award: R01 CA260271. This research used resources of the Argonne Leadership Computing Facility, which is a DOE Office of Science User Facility supported under Contract DE-AC02-06CH11357. The content is solely the responsibility of the authors and does not necessarily represent the official views of the National Institutes of Health. MP was supported by a fellowship from the Belgian American Educational Foundation and a grant from FWO 1161223N. Stanford has submitted a provisional patent application for this work.

## Supplementary information

**Supplementary Figure 1.**
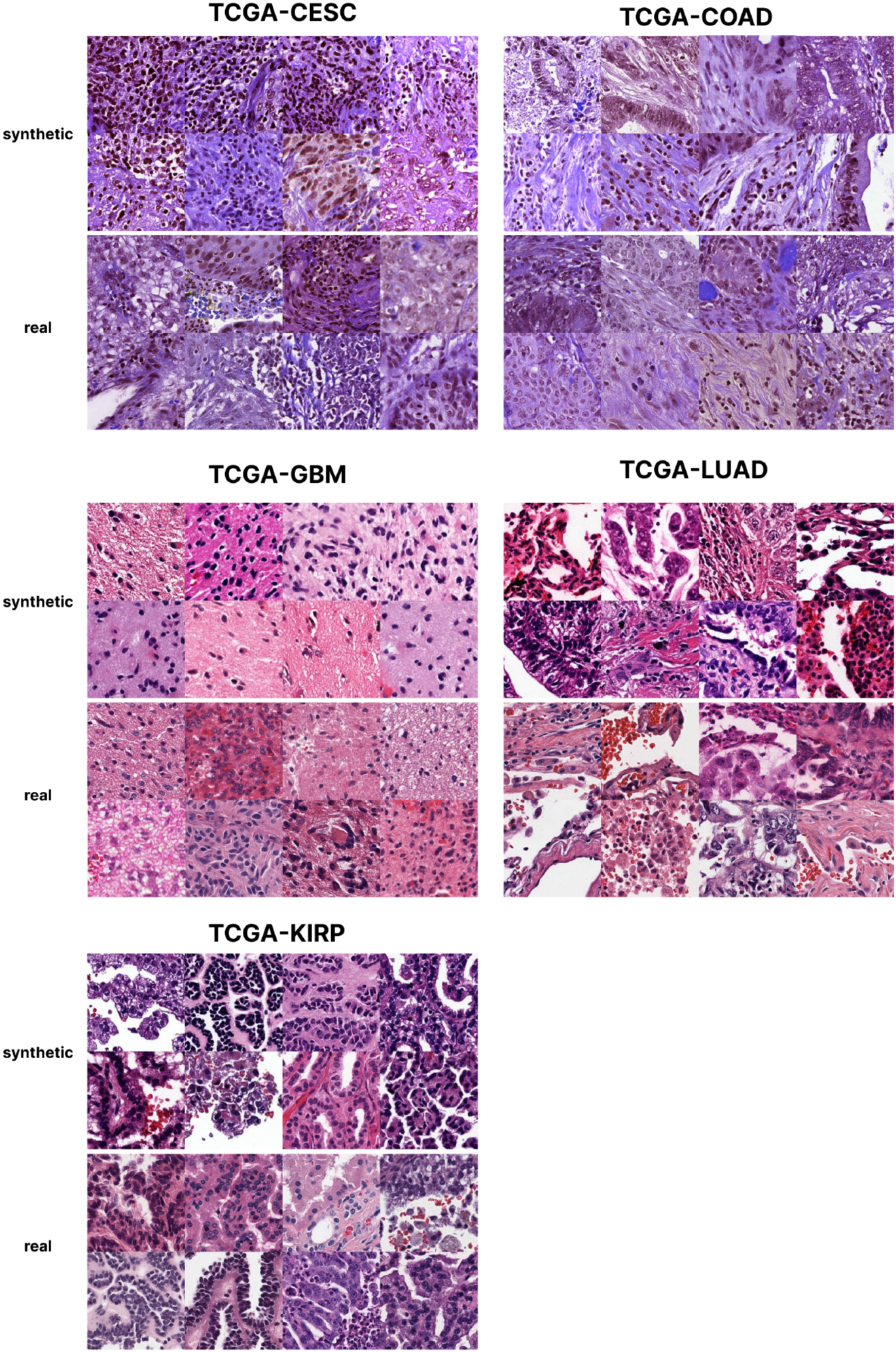
Additional examples of synthetically generated tiles using RNA-CDM compared with real tiles.

**Supplementary Figure 2.**
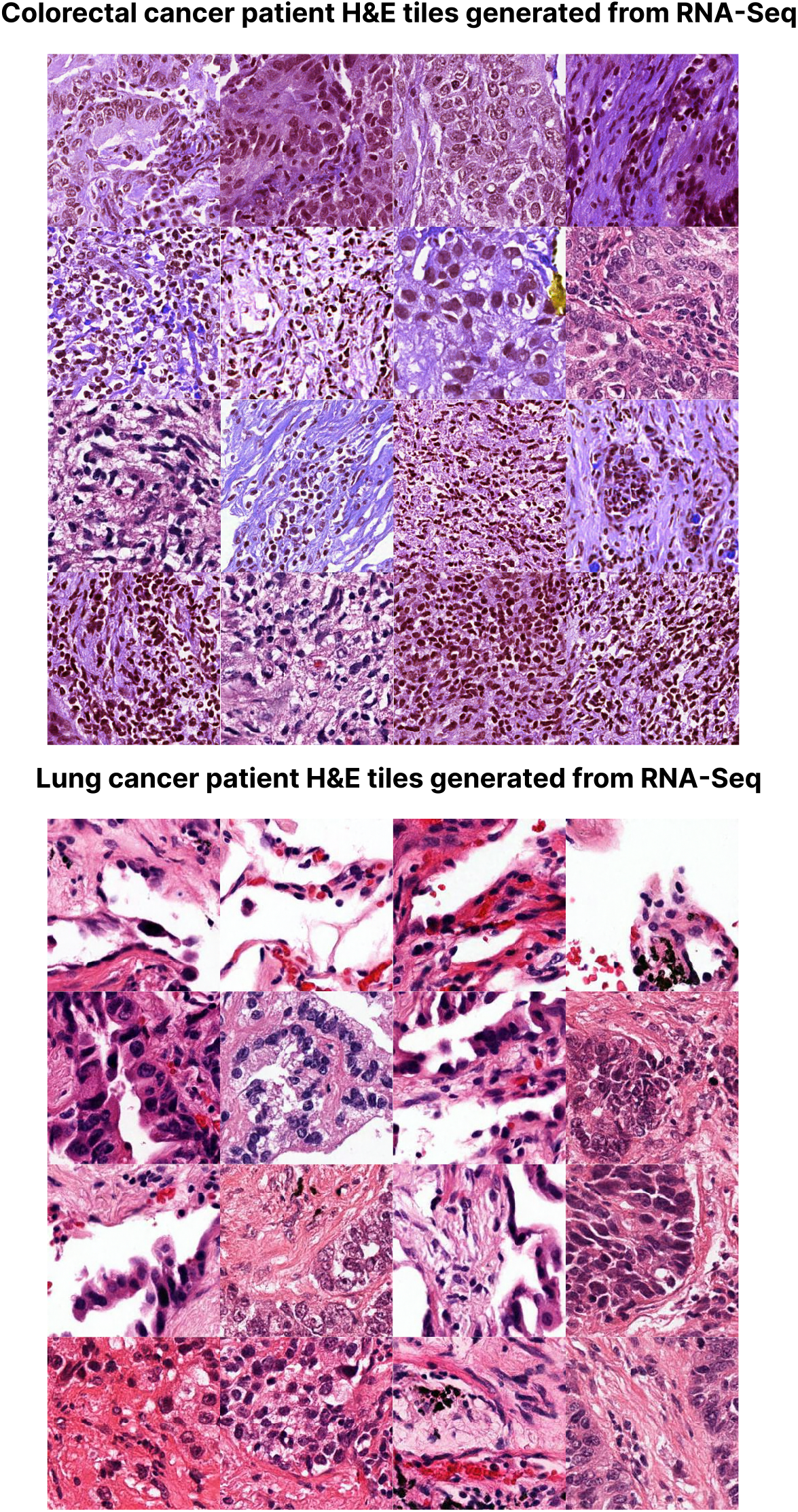
Synthetically generated tiles by using out of the training distribution gene expression from colorectal cancer RNA-Seq [42] and lung cancer RNA-Seq [43].

**Supplementary Figure 3.**
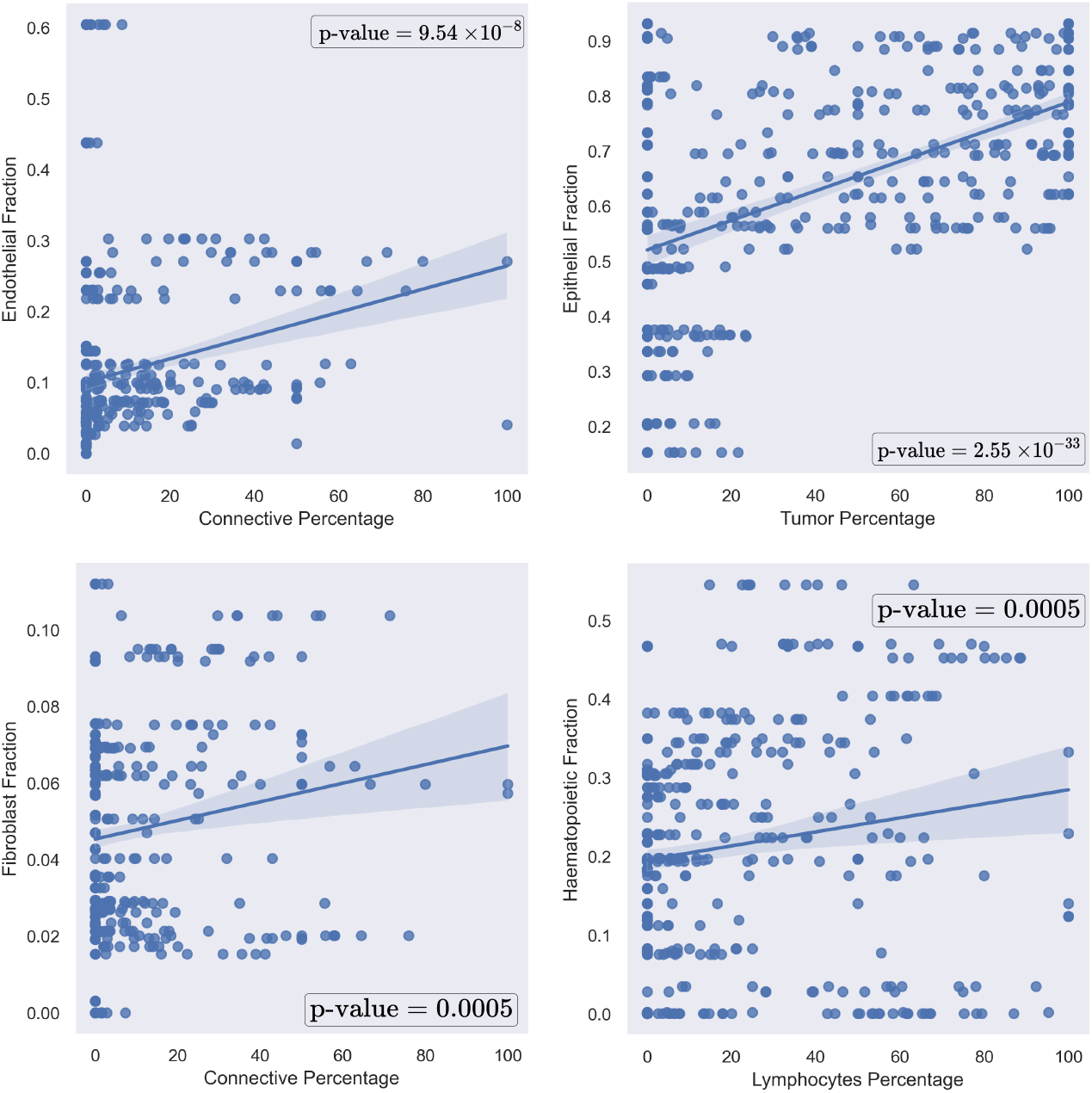
Correlation plots using the pearson correlation coefficient between the fraction percentage of given cells in the deconvoluted RNA-Seq data, and the percentage of cells detected by Hovernet across the different tiles generated using the bulk RNA-Seq data. In all cases there is a significant correlation between them (p-value ≤ 0.05).

**Supplementary Figure 4.**
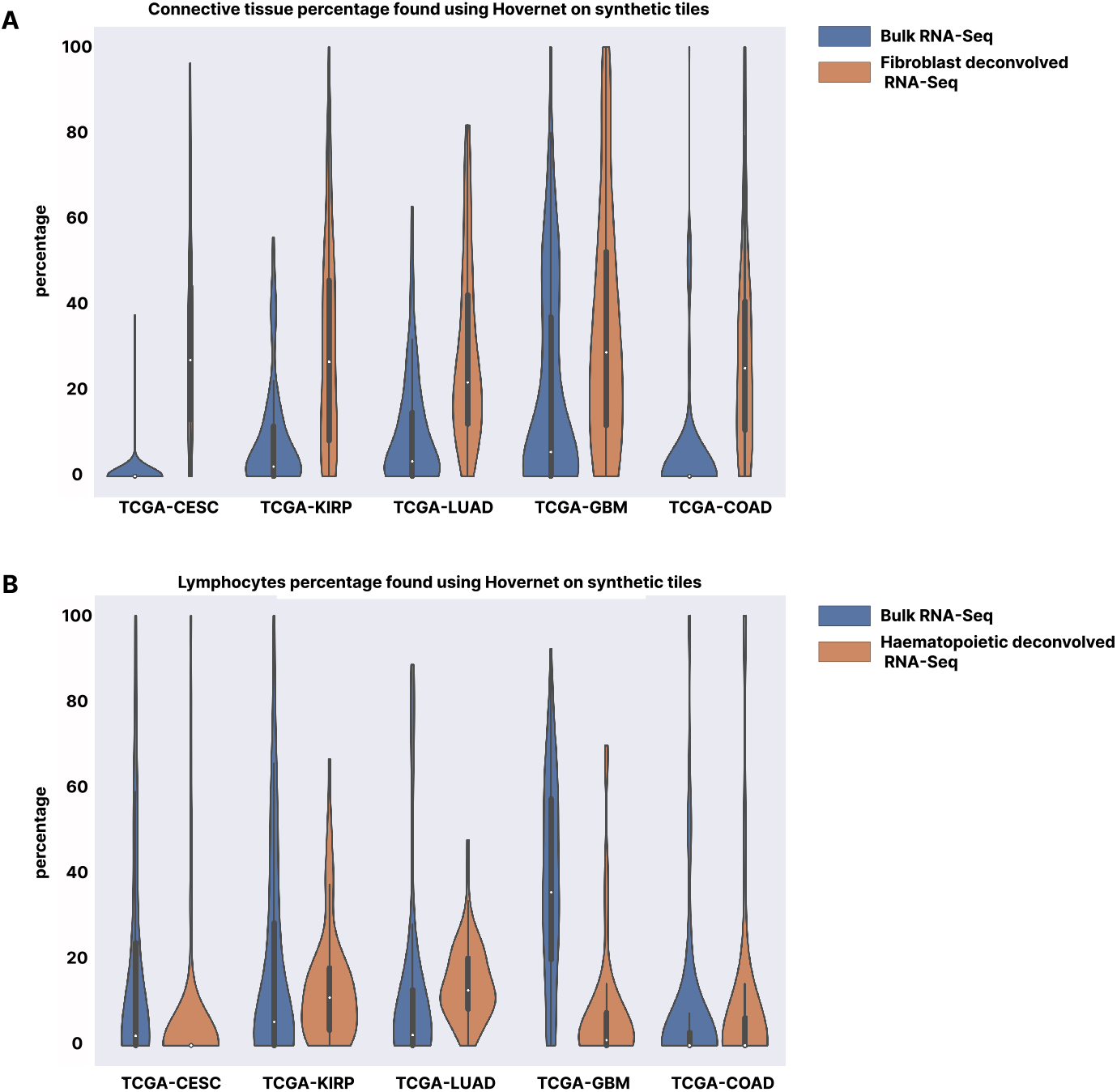
Cell percentage comparison between using bulk RNA-Seq and de-convolved expression. **Panel A:** Percentage of connective tissue cells found by Hovernet in synthetic tiles generated using bulk RNA-Seq and fibroblast de-convolved RNA-Seq. A higher percentage of connective tissue cells are found when the fibroblast deconvolved RNA-Seq across the five cancer types. **Panel B:** Percentage of lymphocytes cells found by Hovernet in synthetic tiles generated using bulk RNA-Seq and haematopoietic de-convolved RNA-Seq. A higher percentage of lymphocytes are found when the haematopoietic de-convolved RNA-Seq in lung cancer and kidney cancer, while a similar amount is maintained in cervical cancer and colorectal cancer.

**Supplementary Figure 5.**
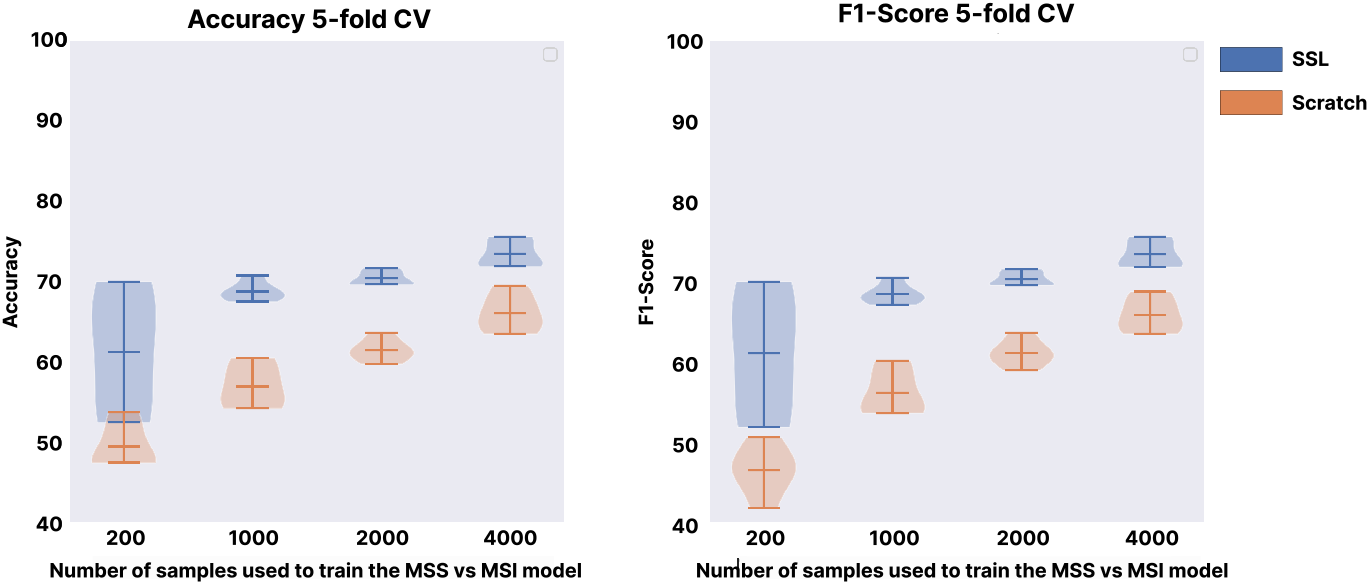
Microsatellite instability status prediction. Comparison between a model trained from scratch and a model that have been pretrained using SimCLR on synthetic tiles, on a different number of real tiles sampled from the training set. Metrics are computed on a 5-Fold CV, and results correspond to those obtained on the different test sets. The model pretrained on the synthetic tiles always outperform the model trained from scratch, no matter the number of training samples that are used.

